# Semiconducting carbon nanotubes decrease neuronal bursting in a network of rat hippocampal neurons *in vitro* while increasing intrinsic excitability of single neurons

**DOI:** 10.1101/2023.03.29.533880

**Authors:** Abhinoy Kishore, Shagnik Chakraborty, Sonali Vasnik, Shinjini Ghosh, Mohammed Raees, Sujit K. Sikdar

**Author notes:** Corresponding Author: Sujit K. Sikdar, Molecular Biophysics Unit, Indian Institute of Science, Bangalore-560012, India, /3220; Email address. The authors contributed equally to the experimental work (Abhinoy Kishore, Shagnik Chakraborty, Sonali Vasnik).

## Abstract

The diverse electrical, chemical and structural properties of the functional derivatives of carbon nanotubes (CNTs) have shown biomedical possibilities for neuroprosthesis or neural interfaces. However, the studies have been generally confined to metallic CNTs that affect cell viability unless chemically functionalized for biocompatibility. Here, we explored the effects of semiconducting single-walled carbon nanotubes (ssw-CNT), on the active electrical properties of dissociated hippocampal neurons *in-vitro* using multielectrode array, calcium imaging and whole-cell patch clamp recordings. The findings show that ssw-CNT treatment regulates neural network excitability from burst to tonic firing by changing the calcium dynamics. However, at a single neuronal level, ssw-CNT increases neuronal excitability.

## INTRODUCTION

Currently, there is interest in studying the integration of nanostructured material with living cells that must be studied for biocompatibility since they affect cellular functioning in diverse ways (Ellis-Behnke et al., 2006; Geller and Fawcett, 2002; Orive et al., 2009; Plant et al., 1997; Silva, 2009, 2006).

Carbon nanotubes (CNTs) are a category of nanomaterials composed of sheets of graphene formed into either single-walled (SWCNT) or multi-walled (MWCNT) carbon nanotubes (Iijima, 1991; Iijima and Ichihashi, 1993) and are investigated as biocompatible scaffolds for neuronal growth for their strength, flexibility, electrical conductivity, and dimensions which are amenable as a substratum for neurons and neuronal structural extensions (Bekyarova et al., 2005; Mattson et al., 2000). The arrangement of sp^2^ hybrid carbon atoms in graphene sheets dictates the conductivity of CNTs (arm-chair conformation-metallic, chiral or zigzag conformation-semiconducting) (Lekawa-Raus et al., 2014; Saifuddin et al., 2012). Neuronal culture on functionalized metallic CNT substrates shows enhanced neuronal activity (Lovat et al., 2005; Mazzatenta et al., 2007), predicting close interaction between neurons and CNTs, and changes in short-term dynamics of synapses leading to spike propagation and active neuronal network (Cellot et al., 2011, 2009), as well as influencing growth by modulating autophagy to facilitate cell attachment (Bekyarova et al., 2005; Xue et al., 2014). However, there are contradictions where nanomaterial-induced cytotoxicity is evident in the form of excitotoxicity and ROS production, structural alterations, activation of molecular pathways (inflammatory and apoptotic), and damages at protein/nucleic acid levels (Fiorito et al., 2018; Jiang et al., 2020) with the susceptibility more prominent in eukaryotic cells (especially mammalian cells) than prokaryotes (Lan et al., 2014). Nevertheless, electro conductive CNTs also show anti-inflammatory and neuroprotective responses by inducing microglial cells, thus regulating CNS activity, structural and functional connectivity, and influencing neuronal plasticity (Fiorito et al., 2018).

Semiconducting CNTs are emerging as an alternative biocompatible nanomaterial to electroconductive CNTs. A study on the comparative and mechanistic toxicogenomics assay of 4 different engineered nanomaterials (ENMs) across three species, viz. *E. coli*, yeast, and human cell lines have demonstrated the IC_5_ (inhibitory concentration at 5%) for metallic and semiconducting CNTs (Lan et al., 2014); while comparing the quantitative levels of toxicity profiles, semiconducting CNTs show a significantly lesser degree of cytotoxicity at similar concentrations of metallic CNTs (Jiang et al., 2020). Metallic CNTs require chemical modification/functionalization to induce biocompatibility. Furthermore, the studies conducted using CNTs involved deposition of the nanomaterial on a substratum and not suspensions in the cell growth medium, limiting their localized depositions in brain structures stereotaxically for functional modifications of neurons for therapeutic purposes. Since, semiconducting single walled CNTs (ssw-CNTs) have not been studied for their effects on neuronal function; here, we explored the effect of ssw-CNT suspensions on dissociated hippocampal neurons *in-vitro* using multielectrode arrays, customized Ca^2+^ two-photon microscopy imaging and whole-cell patch-clamp electrophysiology, to study the changes at neuronal network and single-cell levels.

## RESULTS

### Effect of ssw-CNT on the hippocampal neural population firing properties using microelectrode recordings

The activity of a population of neurons cultured on the MEA (multi-electrode array) dish is a reflection of intrinsic properties of neurons, morphological status of the axon and dendrites that are developmentally regulated as well as the functional communication with other neurons through synaptic activity (Brewer et al., 2009; Kuijlaars et al., 2016; Verstraelen et al., 2018, 2014). We chose to analyze the network activity in vitro with ssw-CNT incubation since it is a good representative for a range of above-mentioned neuronal properties. To determine how ssw-CNTs affect population activity in primary hippocampal neuronal culture, we recorded the spontaneous firing of hippocampal neurons in the presence and absence of ssw-CNTs following different incubation time periods. The timeline of the experiments is shown in Figure 1.

**Figure 1.**
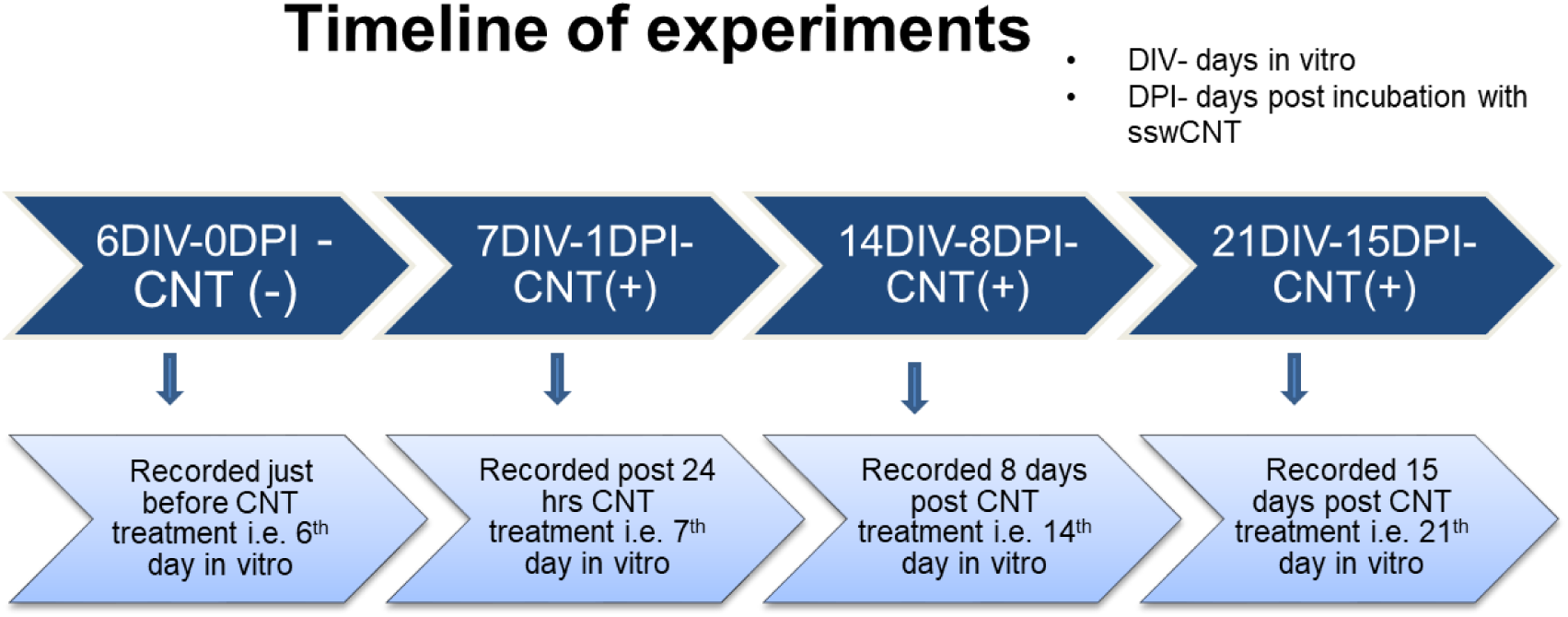
The timeline of treatment of CNT and electrophysiology recordings at specific days of neuronal culture.

The control and ssw-CNT treated neuronal cultures were compared for their baseline neuronal firing properties on 6^th^ DIV just before adding ssw-CNT, to preclude the effect of the initial difference in the properties between the two groups, i.e., baseline mean firing rate, MFR (Figure 2C); number of bursts (Figure 2D); and BFR firing rate, BFR (Figure 2E). This time point is reported as 6DIV-0DPI-CNT (-), where DPI is short for ‘days post-incubation with sswCNTs. There was no significant difference between these two groups on 6^th^ day *in vitro* for MFR (*p* = 0.37590), number of bursts (*p* =0.58546), BFR firing rate (*p* =0.48581). However, neuronal cultures with sswCNTs (SET B) exhibited a decrease in the MFR, number of bursts, and BFR as compared to the control (SET A). This decrease becomes statistically significant at 21DIV-15DPI (MFR: p = 0.03510, number of bursts: p = 0.04580, BFR rate: p = 0.03241; n=10). Furthermore, comparison within SET A at different time points, 6DIV with 7DIV, 14DIV, and 21DIV, also revealed a significant difference on 21DIV in MFR (*p* = 0.00066) and number of bursts (*p* = 0.00068) and significant for both 14DIV (*p* = 0.04439) and 21DIV for BFR rate (*p* = 0.00056). Similarly, for SET B 6DIV comparison with SET B 7DIV, 14DIV and 21DIV showed that the results are statistically significant at 21DIV for MFR.

**Figure 2.**
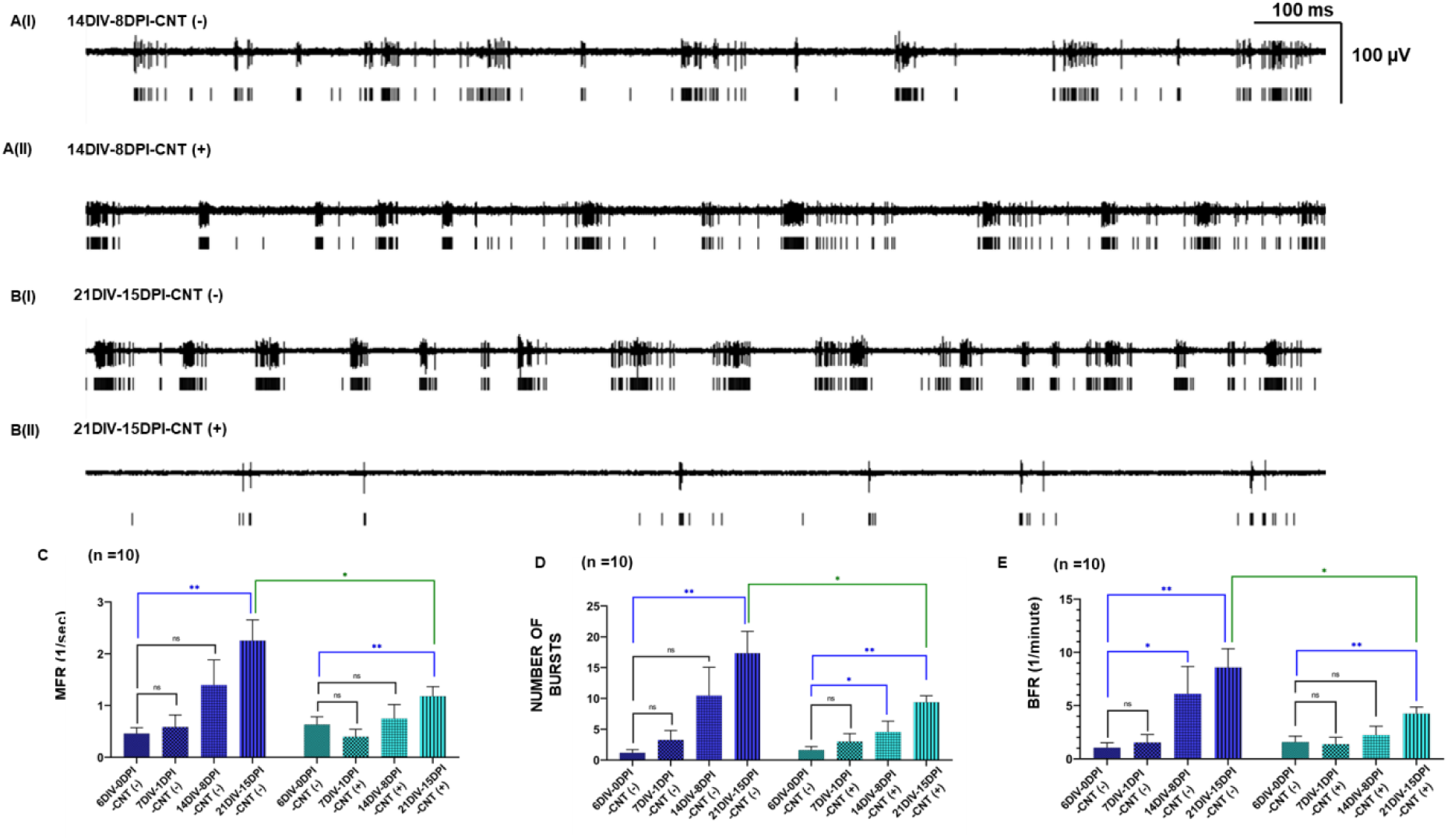
Microelectrode recordings of hippocampal neurons in the presence and absence of ssw-CNTs. Representative traces show the firing pattern of neurons in control (SET A) and ssw-CNT treated (SET B) culture after 14 (AI & BI) and 21 days (AII & BII) *in vitro*. The vertical line below each voltage trace depicts spike timing. An average bar plot comparing mean firing rate (C), number of bursts (D), and BFR rate (E) between two groups. i.e., control and ssw-CNT, and within the same group at 6DIV, 7DIV, 14DIV, and 21DIV *in vitro*; SET A - control, SET B – CNT; n=10 for each experimental group from independent MEAs both in control and with ssw-CNTs.

### Changes in spontaneous Ca^2+^ transients in neurons *in vitro* by ssw-CNT using custom home-built two photon microscopy

To understand the mechanism of decrease in mean network firing dynamics, the intrinsic firing with ssw-CNT incubation was tested at single neuronal level using Ca^2+^ imaging. Carbon nanotubes and films have been reported to cause changes in intracellular Ca ion dynamics in neuronal cells and astrocytes (Jakubek et al., 2009; Ludwig *et al.,* 2020). The effect of ssw-CNT incubation on the intrinsic activity of individual neurons was studied by quantification of Ca^2+^ transients associated with Ca ions influx linked increase in Ca fluorescence due to opening of membrane voltage gated Ca channels during action potential firing by neurons. The shape-based features of the transients viz. Calcium load or the area under the Ca transient curve were measured to understand the changes in intracellular Ca levels during spontaneous neuronal activity in the network culture. Representative examples of neuronal activity as spontaneous Ca^2+^ transients, in control and with ssw-CNT incubation for different time periods are shown in Figure 3. The spontaneous cytosolic oscillations are reminiscent of synchronous Ca oscillations seen in static cultures of hippocampal neurons detected using Ca imaging and patch-clamp electrophysiology, where the Ca oscillations were temporally correlated with bursts of action potentials recorded electrophysiologically in a previous study (Bacci et al., 1999).

**Figure 3.**
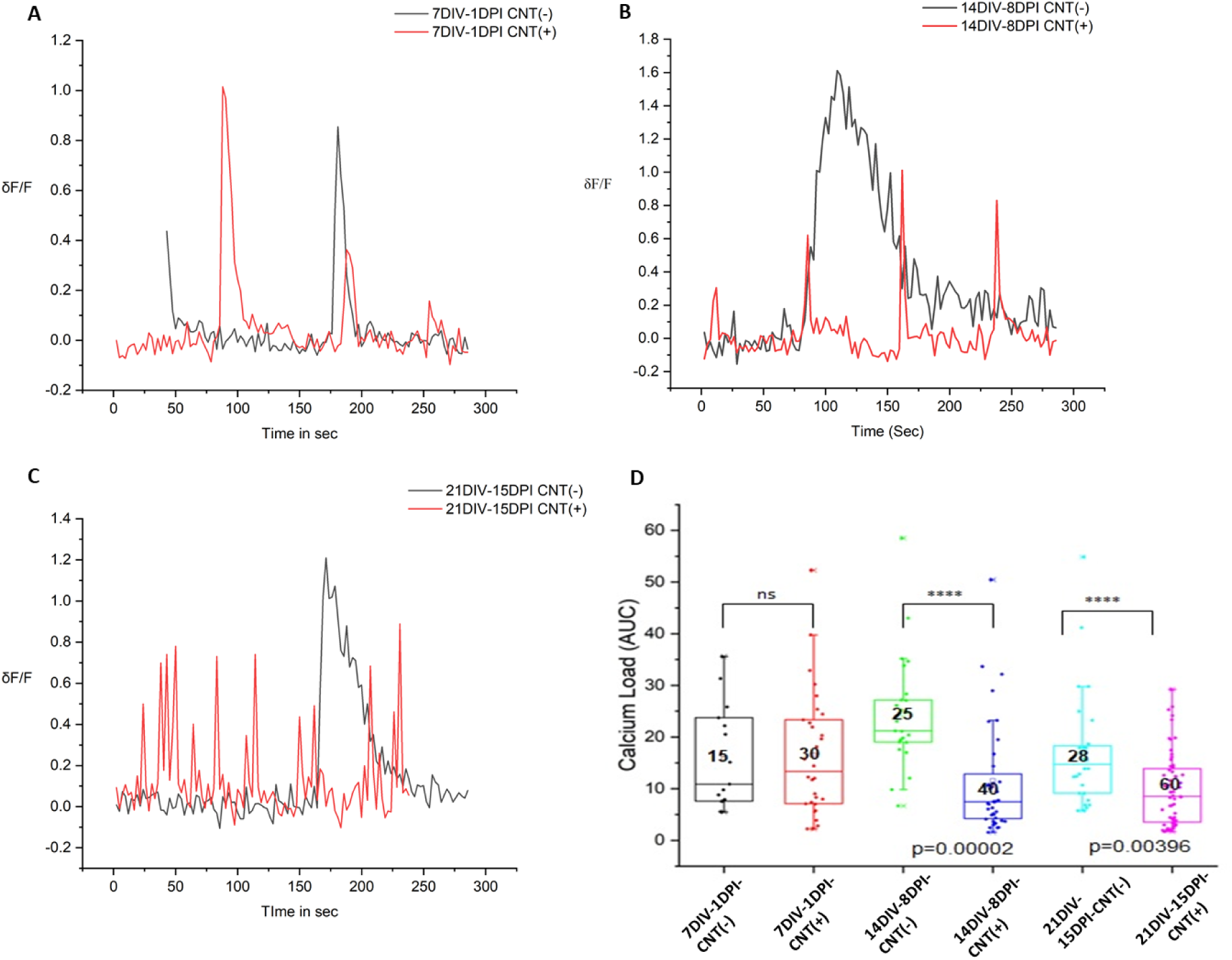
Effect of ssw-CNT incubation on calcium spikes using 2-photon confocal microscopy. Calcium spike at three time periods as the neurons developed in culture in the presence (red trace) and absence (black trace) of ssw-CNT (A-C). The amount of calcium accumulation (represented as area under curve) represented as a box plot at three time periods with and without ssw-CNT treatment (D). The horizontal lines in the box represent medians, box limits indicate the 25th and 75th percentiles, individual data points are represented as open circles and the number inside the box represents the number of neurons (N=3-5 from at least 2 separate cultures) showing calcium transient. Two sample t-test was performed to determine the statistical significance

To understand the mechanism of decrease in mean network firing dynamics, the intrinsic firing with ssw-CNT incubation was tested at single neuronal level using Ca^2+^ imaging. Carbon nanotubes and films have been reported to cause changes in intracellular Ca ion dynamics in neuronal cells and astrocytes (Jakubek et al., 2009; Ludwig et al., 2020). The effect of ssw-CNT incubation on the intrinsic activity of individual neurons was studied by quantification of Ca^2+^ transients associated with Ca ions influx linked increase in Ca fluorescence due to opening of membrane voltage gated Ca channels during action potential firing by neurons. The shape-based features of the transients viz. Calcium load or the area under the Ca transient curve were measured to understand the changes in intracellular Ca levels during spontaneous neuronal activity in the network culture. Representative examples of neuronal activity as spontaneous Ca^2+^ transients, in control and with ssw-CNT incubation for different time periods are shown in Figure 3. The spontaneous cytosolic oscillations are reminiscent of synchronous Ca oscillations seen in static cultures of hippocampal neurons detected using Ca imaging and patch-clamp electrophysiology, where the Ca oscillations were temporally correlated with bursts of action potentials recorded electrophysiologically in a previous study (Bacci et al., 1999).

The ssw-CNT treated neuronal culture showed variations in frequency and amplitude of the Ca transients (Figure 3A-C), while the amplitude of calcium transients and calcium load are decreased in the 14DIV-8DPI CNT and 21DIV-15DPI CNT treated case, these parameters were not affected with 1 day of incubation with ssw-CNTs (Figure 3 B,C). In addition, the burst firing pattern indicated by the long duration Ca transients that have been previously shown to correlate with a burst of action potentials (Teplov et al., 2021) was converted to single sharp calcium spikes after treatment with CNT (Figure 3 A-C). Population data for Ca^2+^ load accord with the results observed in microelectrode recording shown in Figure. 2. Calcium load decreases significantly in 14DIV-8DPI CNT (p=0.00002) and 21DIV-15DPI CNT (p=0.00396) from its control hippocampal culture (Figure 3 D).

### *In vitro* whole cell current-clamped recordings reveal changes in active membrane properties in ssw-CNT treated hippocampal neurons

To further understand the effect of ssw-CNT on the intrinsic firing pattern of single neurons in a culture, changes in neuronal excitability were tested using whole cell current clamp recordings. Incubation of brain slices with carbon nanotubes has been previously reported to enhance neuronal excitability (Varró et al., 2013). However, there are no reports with ssw-CNTs.

*In vitro* patch-clamp recordings revealed that ssw-CNT-treated neurons displayed changes in intrinsic excitability (7DIV-1DPI - Figure 4 A, B and 14DIV-8DPI - Figure 5 A, B). For the 7DIV period, significant changes were observed for P_Amp_ (p = 1.5*10^-4^), and AP_Threshold_, (p = 1.4*10^-7^) (Figure 4 D, E), while the differences in RMP were not significant among the two subsets (Figure 4C); the number of evoked responses (N-eAP vs. I) in ssw-CNT (+) neurons increased for specific current steps with spike adaptation (Figure 4F), while the I_Rheobase_ values significantly differ (p = 6.9*10^-5^) among the two subsets (Figure 4G). Sharper Aps (p = 2.9*10^-8^) were observed in treated subsets quantified using FWHM (Figure 4H), with significantly lesser ΔV (p = 0.02351) value for ssw-CNT (+) neurons (Figure 4I). We observe significantly different values of the MRS (p = 1.4*10^-7^) and MDS (p = 1.3*10^-8^) for ssw-CNT (+) neurons (Figure 4J).

**Figure 4.**
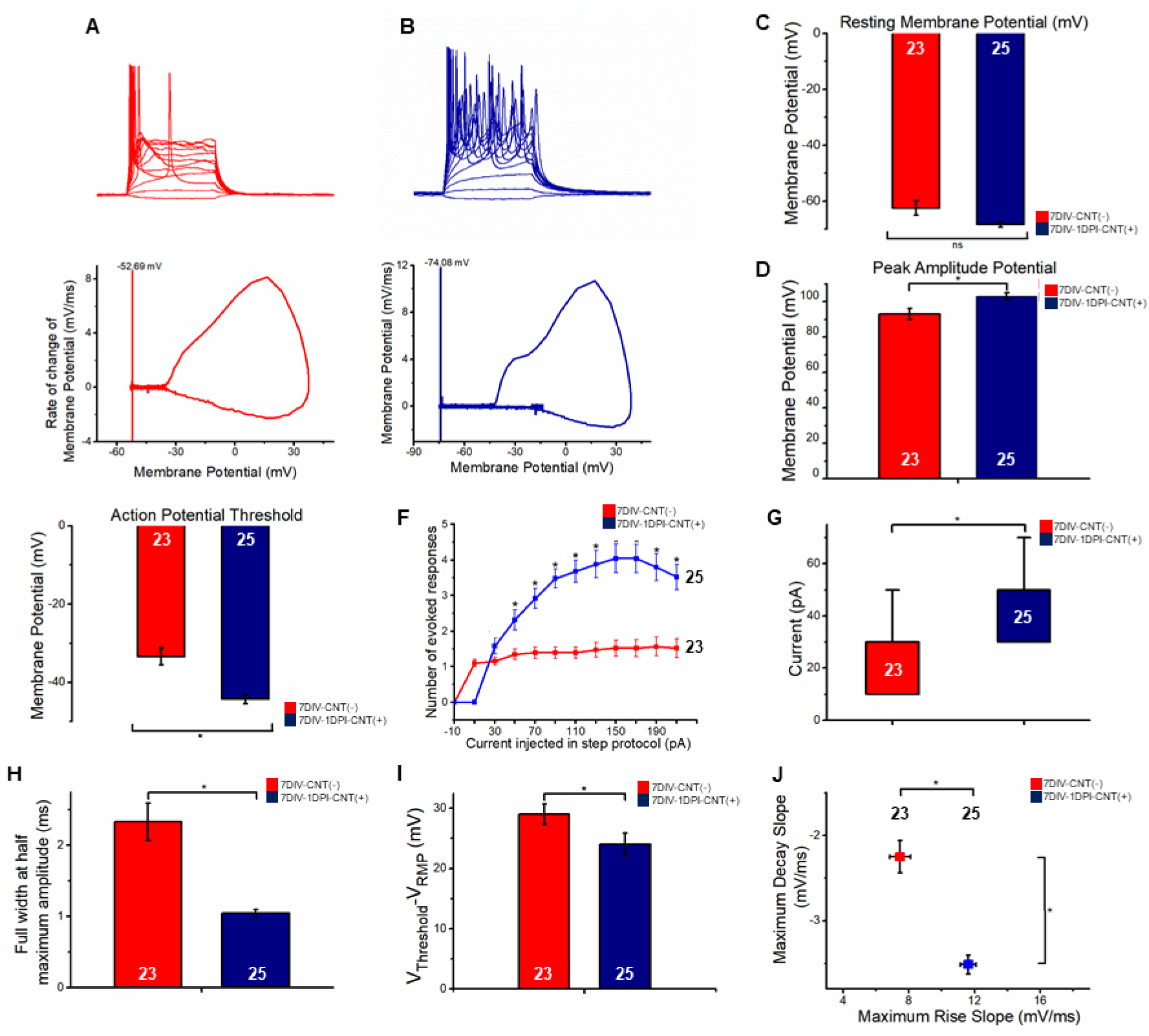
Whole-cell current clamped recordings of hippocampal neurons with absence and presence of ssw-CNT for 7DIV-1DPI time period and the corresponding passive and active membrane properties. Evoked action potential traces devoid of (A) and with ssw-CNT (B) and their corresponding phase plots for the first evoked action potential and showing their respective resting membrane potential. Comparison of passive and active properties of recorded cells (C-I); Resting membrane potential (RMP, C). Peak amplitude potential height (P_Amp_, D). Action potential threshold (AP_Threshold_, E). Number of evoked action potentials vs. current injected at protocol step (N-eAP_Freq_ vs. I, F). near-threshold rheobase current (I_Rheobase_, G). Full width at half maximum (FWHM, H). Responsiveness (ΔV, I). Maximum Rise Slope and Maximum Decay Slope of Action Potential (MRS vs. MDS, J). (Statistical significance has been analyzed by Mann-Whitney test at level of significance of p< 0.05).

**Figure 5.**
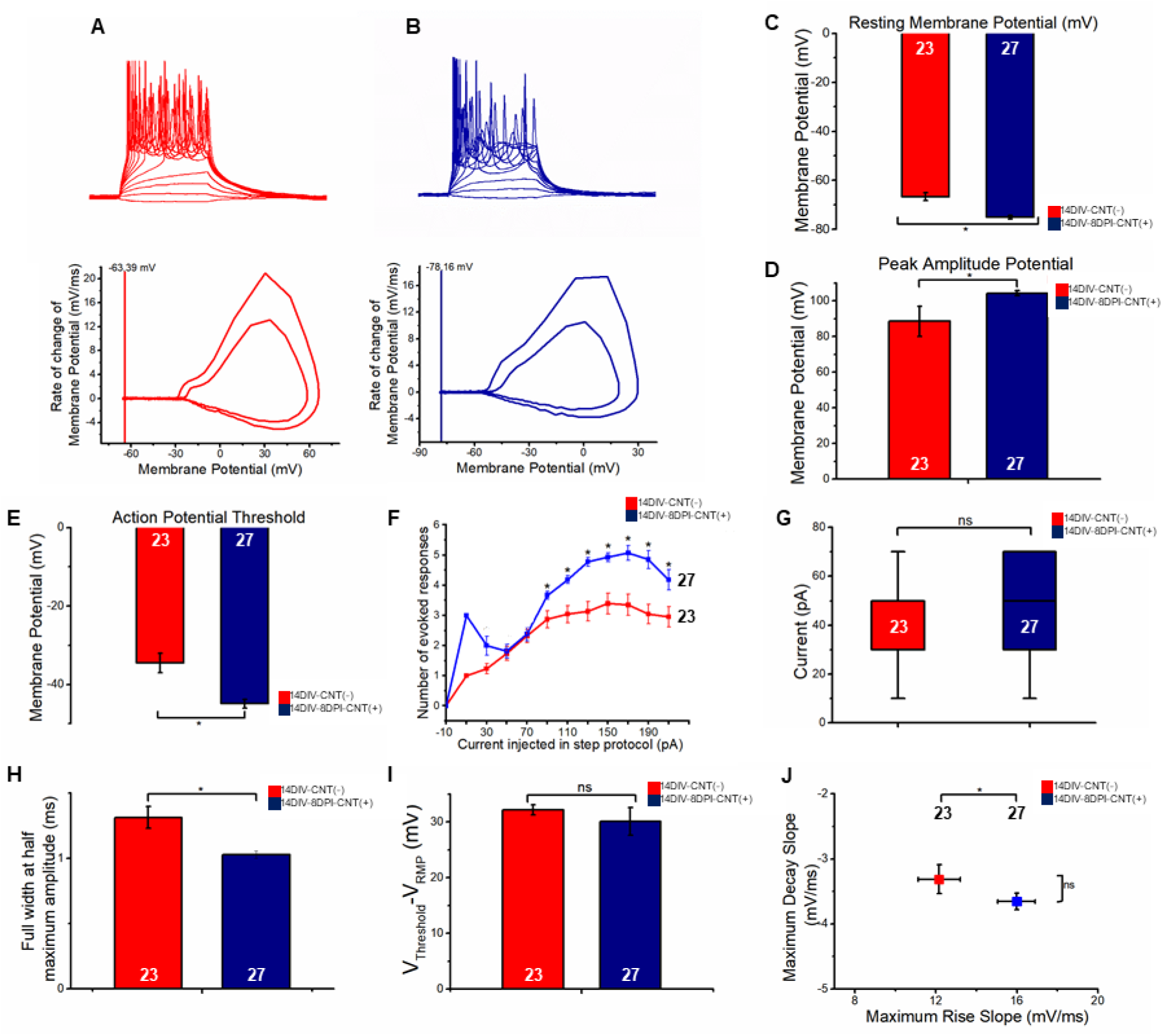
Whole-cell current clamped recordings of hippocampal neurons with absence and presence of ssw-CNT for 14DIV-7DPI time period and the corresponding passive and active membrane properties. Evoked action potential traces devoid of (A) and with ssw-CNT (B) and their corresponding phase plots for the first evoked action potential and showing their respective resting membrane potential (presence of two evoked responses at the rheobase for the representative traces gives rise to two phase plots). Comparison of passive and active properties of recorded cells (C-I); Resting membrane potential (RMP, C). Peak amplitude potential height (P_Amp_, D). Action potential threshold (AP_Threshold_, E). Number of evoked action potentials vs. current injected at protocol step (N-eAP_Freq_ vs. I, F). near-threshold rheobase current (I_Rheobase_, G). Full width at half maximum (FWHM, H). Responsiveness (ΔV, I). Maximum Rise Slope and Maximum Decay Slope of Action Potential (MRS vs. MDS, J). (Statistical significance has been analyzed by Mann-Whitney test at level of significance of p< 0.05).

Contrasting to 7DIV period, for 14DIV period (Figure 5), we see a significant difference between the RMPs of treated neurons to untreated neurons (p = 4.6*10^-9^) (Figure 5C). The P_Amp_ (p = 0.0383) (Figure 5D) and AP_Threshold_ (p = 3.7*10^-4^) (Figure 5E) were significantly different between the two subsets, similar to the 7DIV-1DPI counterpart (Figure 5D) (although untreated neurons showed lower P_Amp_ as compared to the 7DIV-1DPI). Comparing the number of evoked responses at individual current injection steps (N-eAP vs. I), we see spike adaptation in both populations of treated and untreated subsets (Figure 5F), but unlike the 7DIV-1DPI, the I_Rheobase_ values did not differ among the two populations (Figure 5G). Although sharper APs (compared to the 7DIV-1DPI) (p = 3.5*10^-4^) were seen for both subsets, the FWHM for the ssw-CNT (+) neurons is lesser (Figure 5H) while the ΔV values were comparable (Figure 5I). The MRS for the ssw-CNT (+) neurons were significantly higher (p = 0.00763), while the MDS was equivalent for both populations (Figure 5J). Summarizing the changes in passive and active properties of the neuron, we observe sharper APs with a lower threshold, increased amplitude, faster active kinetics, and increased evoked responses for most of the recorded cells for the7DIV-1DPI (Figure 4 C-J). Although the trend in active properties is similar in 14DIV-8DPI, responsiveness, I_Rheobase_ and decay kinetics were not found to be significantly different in the two subsets which suggest that extended incubation do not show further changes to certain active properties (Figure 5 C-J). The changes in the P_Amp_, AP_Threshold_, and MRS and MDS indicate possible alterations in the conduction kinetics/expression of Na_v_ and K_v_ channels respectively (Lam et al., 2017; Zhang, 2004), whose biophysical properties contribute to the measured parameter.

Several studies have shown that CNTs induce changes in neuronal intrinsic excitability; but it is also reported that longer incubation with CNTs causes increased oxidative stress related toxicity on neurons in culture. On the contrary, our experimental results show ssw-CNTs may not induce significant oxidative toxicity on neurons as reported to occur for other categories of CNTs since neurons showed electrical excitability even after 16 day incubation with ssw-CNTs and following 27 day incubation as well (data not shown) although quantitative assays using biomarkers specific for neuronal viability would further substantiate the observations (Jiang et al., 2020).

## DISCUSSION

Carbon-based Nanomaterials (CBNs), especially carbon nanotubes (CNTs) are the center of research for prospective materials for neuroprosthesis. But the observed toxicity is a deterrent for devising therapeutic strategies for neuronal connectivity restoration. The investigation of the effects of CBNs on the electrical activity of neurons mostly mentions metallic CNTs (Cellot et al., 2011, 2009; Lovat et al., 2005; Mazzatenta et al., 2007); the absence of an energy gap between conduction and valence band might stimulate a neuron continually (unlike in-vivo conditions) by functioning as a conductive bridge to couple the somatic and dendritic compartments (Cellot et al., 2009) which could be detrimental for neuronal viability in the long run. Semiconducting single walled CNTs (ssw-CNTs) used in the present work barely induce ROS production and genotoxicity in neurons (Jiang et al., 2020) compared to metallic CNTs.

The normality of the neural network development in our control set of cultures is reflected in the spontaneous firing tendency of neurons. A set of studies have reported the developmental changes of neuronal culture on MEA, wherein the 1^st^ week the neurons fire randomly but in the 2-3^rd^ week the synaptic connections become organized such that the network fire more synchronously in bursts (Cotterill et al., 2016; Li et al., 2005; Negri et al., 2020). As seen in Figure 2C, our neuronal cultures on MEA dishes showed normal course of development with a progressive increase in the mean firing rate from 0DPI to 14DPI in culture showing a sigmoidal trend signifying the functional maturity of the neuronal network with the formation of synaptic connections leading to progressively increase in the activity. Interestingly, while the trend of progressive enhancement in firing characteristics of the network with development indicated by the increase in MFR remained with ssw-CNT incubation, this was tuned down (Figure 2D). A significant decrease in the mean firing rate post 14-day CNT treatment suggests the neuromodulatory role of ssw-CNT on the network property of hippocampal neurons in primary culture.

Our neuronal network cultures *in vitro* showed burst firing. During bursting activity of neurons, alternate bouts of quiescence when there is no firing of action potentials or sparse firing, and repetitive neuronal firing is observed. Neuronal cultures *in vitro* have been previously reported to show bursting activity as they develop in the culture as part of the developmental process associated with extension and maturation of synaptic contacts (Lillis et al., 2015; Wagenaar et al., 2006). The mechanisms responsible for burst generation include both intrinsic properties of a single neuron or as a local network activity involving synaptic transmission and both mechanisms can interact to generate bursts. Neurons that generate bursts in isolation are called intrinsic bursters. This usually requires the participation of a T-type voltage gated Ca channels (Lambert et al., 2014) to induce slow membrane depolarization on top of which repetitive action potentials are generated. Neurons that are not intrinsic bursters in a network can still generate bursts of action potentials as an emergent property of the network due to synaptic mechanisms.

Since the bursts characterized in our MEA recordings were from neurons that are part of a network, the burst recorded from neurons in the MEA (Figure 2) could be due to the combined contribution of intrinsic mechanisms and synaptic mechanisms in the network. A significant observation was the reduction in the burst characteristics and numbers with ssw-CNT treatment (Figure 2E).

While the mechanisms underlying the decreased network bursts are difficult to propose at this stage, this could in part be explained by the changes in intracellular free Ca levels based on the findings of our Ca fluorescence imaging studies where broad Ca transients representing bursts of action potential firing in control changed to more sharp Ca transients (Figure. 3). Calcium fluorescence transients reflect intracellular calcium level dynamics and encode the action potential frequency and pattern. High-frequency burst patterns of 100-200 Hz are not resolved in Ca^2+^signal and appear as broad spikes (Helmchen et al., 1996). Different algorithms exist that transform calcium signals into trains of action potentials (Quan et al., 2010).

The acquisition rate (frame interval of 2.38s,) was similar in both control and treated conditions, thus the transition of broader calcium transient peak in control set to sharp spikes would predominantly reflect the ssw-CNT induced changes that cannot be entirely accounted by the slow acquisition frame rate alone. Following 7DPI with ssw-CNTs a significant decrease in the peak amplitude of the Ca transients and the Ca load were observed. Previous study by (Bacci et al., 1999) have clearly shown that synaptic and intrinsic conductances shape the synchronized Ca oscillations in hippocampal neurons in static cultures, similar to ours. The simultaneous activation of of N-methyl-D-aspartate (NMDA) receptors due to glutamatergic synaptic activity and opening of voltage-activated L-type calcium channels allows calcium influx that regulates the spontaneous Ca transients (Bacci et al., 1999) It is noteworthy to mention here that multiwalled carbon nanotubes have been previously reported to inhibit glutamatergic neurotransmission, although the cellular mechanisms involved were not clear (Chen et al., 2014). Since the Ca transients are associated with the opening of voltage-gated Ca ion channels on the neuronal membrane that are activated during action potentials, decreased Ca influx could in part be explained by the block of voltage-gated Ca ion channels by ssw-CNTs. In fact, the block of voltage-gated Ca ion channels by CNTs has been reported earlier (Jakubek et al., 2009). An additional increase in the intracellular Ca buffering indicated by the faster decay of the Ca transients by ssw-CNTs through upregulation of Ca buffer proteins of intracellular Ca is possible.

We postulate that ssw-CNTs may influence the bursting behavior of the neurons in the network by partial block of T-type Ca channels and decreasing the release of excitatory neurotransmitters that are dependent on intracellular Ca levels. In addition, ssw-CNTs may influence neuronal network connectivity in cultures, going by the interesting experimental observations of (Ivenshitz and Segal, 2010) where neurons in sparse cultures that were experimentally achieved by controlling the density of neurons while culturing, showed spontaneous neuronal activity with enhanced burst size but reduced burst frequency compared to dense cultures. We postulate that ssw-CNTs can increase neuronal connectivity by enhancing synaptic contacts with neighboring neurons.

Although we are unable to provide the mechanism underlying the interaction of our nanomaterial with the neurons, there is a possibility of non-toxic concentrations of ssw-CNTs translocating across the cell membrane because of its dispersibility or getting lodged across the lipid bilayer (of neurons, astrocytes and/or microglia) to function as selective solid-state ionic channels and activate downstream biochemical cascades that show cytoprotective effects in the cellular subtypes (Fiorito et al., 2018; Jiang et al., 2020; Xue et al., 2014). Neuronal intrinsic excitability could be modulated through mechanical/electrical interactions across smaller diameter neuronal bundles which influence channel gating (Cellot et al., 2009), or the formation of neuronal “short-circuits” only upon stimulation which limits the excitation-mediated toxicity. Previously, Pampaloni et al. (2018) showed the π electron clouds of sp^2^ carbon networks single layer graphene (SLG) interferes with ion bulk diffusion by adsorbing the ions to tip the ionic driving force to change cellular excitability; using a minimal *in-silico* model it describes how SLGs increase the intrinsic firing of neurons by modulating ion conduction at the graphene-neuron interface; hypothesizing from the same we assume that the spike-adapting excitatory responses by ssw-CNT could be due to possible modulation of Na^+^ conductance as observed from the distinct features in the evoked responses of the treated neurons (Figure 4 and 5). Additionally, reduced intracellular Ca following ssw-CNT incubation (Figure. 3) can also in part account for the enhanced activity of neurons by ssw-CNTs through the reduction of Ca-dependent K ion channel activity that normally functions to reduce neuronal firing (Vierra and Trimmer, 2022). Thus, molecular dynamics simulations of the interactions of the triad of nanomaterial, membrane ion channels and specific ions (Na^+^, K^+^, Ca^2+^, Cl^-^) might give us clues about a possible mechanism of altered ion channel kinetics.

The systematic analyses conducted using ssw-CNT showed promising outcomes to function not only as a prospective nanomaterial for developing neuro-hybrid systems, to create tunable band-gap constructs that sense thermal energy, physical interaction or extracellular field potentials. This modulates neuronal activity and neuronal morphology at the level of dendritic spines that are regions of synaptic contacts, but that could also be used for neurotherapeutics (Cellot et al., 2011, 2009; Mazzatenta et al., 2007; Serpell et al., 2016).

The decrease in the bursting activity of neurons by the ssw-CNTs (Figure. 2) potentially has a direct neurotherapeutic application in neurological diseases like Parkinson’s, where the motor dysfunction is associated with increased bursting activity in discrete areas of the brain such as the basal ganglia nuclei (Lobb, 2014). Stereotaxic local delivery of ssw-CNTs at the affected regions could be used to ameliorate the condition by reducing the bursting activity to normal non-pathological levels to restore motor dysfunction. Depression is another mental condition of mood disorder, associated with increased bursting in the lateral habenula (Shao et al., 2021) where ssw-CNT could be used. A miniaturized neural drug delivery system recently developed (Dagdeviren et al., 2018), could be used for localized delivery of the ssw-CNTs in the brain.

In conclusion, the promising results of our investigation suggest a nanomaterial with a tunable energy band-gap for CNT-based neuronal interfaces, implants in damaged tissues in the form of scaffold/suspension, or as virus-based targeted nanoparticle delivery methods (Koudelka et al., 2015) to modulate structural and functional connectivities between neurons.

## EXPERIMENTAL DESIGN

Inclusion or exclusion criteria of any data or subjects: We did not exclude any data from the analysis. All data presented in the figures were biological replicates as they were from separate samples and are listed in the figures as sample size.

## MATERIAL AND METHODS

### Primary hippocampal neurons culture and treatment of ssw-CNT

Hippocampal neuronal cultures were obtained as follows (Rao and Sikdar 2004; Srinivas et al. 2007): Briefly, hippocampi isolated from p0-p2 Wistar rat pups were treated with papain solution (20 U/mL enzyme activity at 25°C) for 30 min and then washed with culture medium. They were then mechanically dissociated by trituration and filtered through a nylon mesh. The filtered homogenate was spun in a centrifuge at 1000g for 5 min; the supernatant was discarded, and the pellet was resuspended in culture medium. The cell suspension was then seeded directly on PDL coated 35 mm coverslips and MEA dishes (120MEA200/30iR-Ti Multichannel Systems) at cell density of 1×10^3^ cells/mm^3^ and maintained in a humidified incubator supplied with 5% CO_2_ at 37°C One-half of the culture medium was changed every 3 days. The culture medium contained DMEM F12 HAM with 1% N-1 supplement, 10% FBS and 1% antibiotic-antimycotic. 10 μM Cytosine β-D arabinofuranoside (Ara-C) was added after few days in culture to limit astrocyte overgrowth. The studies were restricted to the following time periods viz. 7 (7DIV-1DPI), 14 (14DIV-8DPI) and 21 (21DIV-15DPI)-days post-culturing of neurons. Semiconducting Single walled carbon nanotube stock was diluted 10 times, sonicated for 20 minutes and then added (1µg/ml) over the matched neuronal culture subsets of the batch culture on 6th day in culture. Both short-term, where the neuronal cultures were incubated with sswCNT for 1 day, and long term, where the neuronal cultures were treated with sswCNT for more than 7 days were studied for their effects on neurons. The neurons were thus allowed to grow in the presence of sswCNT for short and long duration. Changes induced by ssw-CNT were compared with matched control from the same neuronal culture batch.

### Multielectrode Arrays recordings

The MEA-2100 System from MultiChannel Systems, Germany© was used for recording from the dissociated neuronal cultures on the MEA. The system consists of a head-stage and an interface board. The ADC (Analog to Digital Converter) headstage can sample at the rate of 50 kHz on 120 electrodes. This provides sufficient resolution for acquiring data from neurons which is suitable for spike-detection. We used MC_Rack software from Multi Channel Systems, Germany for recording.

The recorded files (.mcd) were loaded in the Neuroexplorer (Plexon v5). The spikes were extracted using amplitude thresholding after bandpass filtering the signal (300 Hz to 6000Hz, four pole Butterworth filter). The standard deviation (SD) and the median of the signal absolute values were calculated (MA), from which the median sigma (MS) was determined by MA/0.6745. The custom threshold was set to negative threshold of 5 times the SD of the baseline noise and spikes that crossed MS * custom thresholds were detected. Burst detection was done using the MaxInterval method. The parameters like Max ISI in a burst, Interburst interval, and minimum burst duration were determined by logISI plots and was found to be 20 ms, 0.2 s and 20 ms respectively while other parameters like MaxISI at start of the burst (0.2s) and number of spikes in a burst (3) were taken from standardized values from (Cotterill et al., 2016). From here, the BFR (1/minute), MFR (1/s), and a total number of bursts across each electrode were calculated.

#### 2-Photon Calcium ion transient imaging using Calcium Dye-Cal 520 AM

The Two Photon microscope system built in-house had a Titanium Sapphire laser (Chameleon vision II) which produced 70fs pulses with wavelength tunability ranging from 680-1060nm with an average power >1W and beam diameter of 1mm. A small fraction of the beam was reflected to the spectrum analyzer (USB4000, Ocean Optics) for checking the spectral bandwidth. The laser beam was expanded by a Keplarian based telescope to a diameter of 2.5 mm and intensity control was achieved by half waveplate and polarizing beam splitter (Thor Lab Inc., USA) combination. A pinhole placed at the focal plane between the telescopes lenses served as a spatial filter to provide TEM 00 beam. An electronic high-speed shutter (Uniblitz VCM-D1 shutter controller, USA) blocked the laser beam when no images were taken. A Neutral density filter (Thor Lab Inc., USA) wheel was used to reduce beam intensity further. Working with hippocampal slices, the beam intensity was adjusted to 8-15mW of average power onto the object plane without causing any detectable changes in the cell electrical properties. The IR beam was guided by mirrors to the laser coupling port of the scan head of the Zeiss AxioExaminer Z1 microscope. A mirror reflector then directed the scanning beam down to the microscope. In most of the experiments, a Zeiss AplanoChromat 20X, 1.0 NA, water immersion objective with long working distance was used. The scanner system was equipped with a dual galvanometer (GVS0002, ThorLabs Inc., USA) scan head and drive electronics. The two galvanometer mirrors separated by 10mm were mounted on a large scan unit. The galvano-mirrors were controlled by a saw tooth voltage command waveform generated by data acquisition system (PCI 6110, National Instrument). The hinge point sits in between the two scanning mirrors which is relayed back to the focal plane of the 20X objective with the help of an intermediate telescope consisting of the scan lens and the tube lens. Thus, the change of the deflection angle by the scanning mirrors was translated into a lateral movement of the focal spot. To obtain diffraction limited optical resolution we had to overfill the back focal plane of the objective. In the back focal plane of the objective, the laser beam was collimated without much lateral beam drift following the movement of the scan mirror. An excitation dichroic mirror (FF 705-Di-01, Chroma) just above the objective was used to separate the IR excitation light from the fluorescence light. The fluorescence was then reflected to the dual channel detectors with emission filters. Fluorescence detection was done in the wide field, non-descanned detection mode using photomultiplier tubes (PMT H7422-40, Hamamatsu, Japan). This detection scheme has high photon collection efficiency. On the detector side stray IR light was blocked by a BG-39 filter (Thorlabs, USA). Fluorescence light was split for each detector pair by a dichroic (FF 568-Di-01, Chroma) separating green from the red channel. Band pass filters (Green, ET535/30m, Red, ET630/75m) in front of each detector were employed to prevent cross contamination. The demagnification telescope (5:1) in the detection side enables light collection within the available 5mm photosensitive area of PMT.

Cal520 acetoxymethyl ester (Cal-520 AM) dye was prepared according to the protocol defined by the manufacturer. Briefly, solution of 20% Pluronic F-127 in DMSO was freshly prepared and sonicated for 10 min. Four microliters of this solution was used to reconstitute 50 µg of powdered Cal-520. The resulting solution was sonicated for another 12–15 min and then brought to a total volume of 40 µL by adding 36 µL of a calcium-free solution (in mM: 150 NaCl, 2.5 KCl, and 10 HEPES, pH 7.4) to reach the final concentration of 1.13 mM Cal520-AM. The working solution was prepared by diluting 2.5 µL of the above solution to 500 µL by calcium free solution. The dissociated primary hippocampal neuronal culture was incubated with a working solution of Cal520 AM for 20 mins at 37 °C. After 20 mins, the solution was removed and washed thrice with an External solution. To obtain calcium transients from neuronal populations, images were acquired at a resolution of 512×512 pixels for 2 min at a frame rate of 0.42HZ (i.e., 2.38sec per frame) to generate the image stack and analyzed using Fiji-ImageJ software for previously defined time periods viz. 7DIV-1DPI, 14DIV-8DPI and 21DIV-15DPI.

Total of 120 images were acquired in each stack. The image stack was analyzed using Fiji-ImageJ software. The region of interest (ROI) was selected in active neurons and the average of fluorescence signals was calculated.

Calcium transient was quantified as δF/F = (F-F_0_)/ (F_0_-F_b_), where F_0_ is baseline fluorescence and F_b_ is background fluorescence for Cal520-AM.

The activity of individual neurons was monitored by studying changes in intracellular Ca ++ fluorescence that are linked to opening of voltage gated Ca channels. At rest, in general the concentration of Ca ion is 50-100nM and this can rise transiently during electrical activity by10-100 times. The dissociated hippocampal culture was grown in presence of semiconducting single walled carbon nanotube (sswCNT) for various time points. The Ca^2+^ transient, representative of neuronal activation, was captured for series of days *in vitro* (DIVs).

#### In-vitro Electrophysiology

Whole cell patch clamp electrophysiology data acquisition and analysis

Whole cell recordings were performed at room temperature on neurons possessing oval or pyramidal-shaped somas, which were observed under the 20X objective lens of the Axio Observer Z1 Inverted Microscope (Carl Zeiss). All recordings were performed using a Multiclamp 700B amplifier (Molecular Devices) with a Digidata 1440A analog-to-digital converter (Molecular Devices), filtered at 2 kHz and digitized at 10 kHz. Current clamp protocol was applied using WinWCP Whole Cell Analysis Program v5.4.1 (Strathclyde Electrophysiology Software).

Pipettes with resistance of 3.5-6 MΩ were fabricated from borosilicate filamented capillaries using the P-97 micropipette puller (Sutter Instrument). The pipettes were filled with an internal solution composed of 8mM NaCl, 0.6 mM MgCl_2_•6H_2_O, 0.1 mM CaCl_2_•2H_2_O, 125 mM K-gluconate, 10 mM HEPES, 1 mM EGTA, 4 mM Mg-ATP, and 0.4 mM Na-GTP. The micro cover glasses, with the attached cells, was transferred to the recording chamber and was bathed in an external solution comprising of 125 mM NaCl, 1 mM MgCl_2_•6H_2_O, 3 mM CaCl_2_•2H_2_O, 2.5 mM KCl, 10 mM HEPES, and 20 mM glucose, and its osmolarity was adjusted to 305–310 mOsm with sucrose. The osmolarity of the internal solution was always kept at a value 10 mOsm lower than that of the external solution. The pH of both the internal and external solutions was kept between 7.2-7.4 (Verma et. al. 2017).

The pipette offset and series resistances were compensated beforehand. The current clamp recordings involved step pulses of 20pA with an initiation value of -10pA for a duration of 150ms. We divided our electrophysiology studies into two time periods: 7DIV-1DPI (7days) and 14DIV-7DPI (14days) to study the changes in the electrical properties of neurons devoid of semiconducting-single-walled CNTs (ssw-CNT) and with ssw-CNT for 1 and 8 days; the recordings were restricted to the above periods only due to the excessive coverage of astroglia post 19 DIV which made neurons less accessible for proper patching (Grabrucker et al., 2009). The current clamped studies enabled us to represent the nature of the evoked responses and corresponding phase plots for the near-threshold rheobase current injection step, and determine the following parameters of Action Potential (AP) viz. Resting Membrane Potential (RMP), Peak Amplitude Potential (P_Amp_), Action Potential Threshold (AP_Threshold_), number of evoked action potentials vs. current injected at protocol step (N-eAP vs. I), near-threshold rheobase current (I_Rheobase_), Difference between Threshold Potential and Resting Membrane Potential (ΔV), Full Width at Half Maximum (FWHM) and Maximum Rise Slope vs. Maximum Decay Slope for an Action Potential (MRS vs. MDS) were determined for the different sets in the aforesaid time periods. Recording data visualization, quantification and analysis were done using Clampfit 10.6 software (Molecular Devices) and OriginPro 9.0.0 SR2b87 (OriginLab Corporation). The associated threshold values were determined by matching the first peak of the third derivative with the original trace and the point of intersection was noted as the threshold (Henze and Buzsáki, 2001). The peak action potential amplitude height was measured as a difference between the resting membrane potential of the cell and the peak membrane potential value under evoked responses. The comparison of the number of evoked action potentials at each current injection step was quantified by the total number of action potentials that were observed in one current injection sweep of the current clamp protocol and measured similarly for the entire protocol sweep. The action potential maximum rise and decay slopes, and phase plots were quantified using the rheobase step elicited by our current clamp recording protocol. The FWHM was calculated at the halfway point of the amplitude using Python 3.7.4. The corresponding active and passive properties of the recorded neurons were utilized for Principal Component Analysis for dimensionality reduction and to discover potential relationships among the variables and to derive conclusions about possible systematic structures among the two populations in the two time periods; each dot represents the projection of a quantifiable feature of the biophysical properties of the neuron.

The pipette offset and series resistances were compensated beforehand. The current clamp recordings involved step pulses of 20pA with an initiation value of -10pA for a duration of 150ms. We divided our electrophysiology studies into two time periods: 7DIV-1DPI (7days) and 14DIV-8DPI (14days) to study the changes in the electrical properties of neurons devoid of semiconducting-single-walled CNTs (ssw-CNT) and with ssw-CNT for 1 and 8 days; the recordings were restricted to the above periods only due to the excessive coverage of astroglia post 19 DIV which made neurons less accessible for proper patching (Grabrucker et al., 2009). The current clamped studies enabled us to represent the nature of the evoked responses and corresponding phase plots for the near-threshold rheobase current injection step, and determine the following parameters of Action Potential (AP) viz. Resting Membrane Potential (RMP), Peak Amplitude Potential (P_Amp_), Action Potential Threshold (AP_Threshold_), number of evoked action potentials vs. current injected at protocol step (N-eAP vs. I), near-threshold rheobase current (I_Rheobase_), Difference between Threshold Potential and Resting Membrane Potential (ΔV), Full Width at Half Maximum (FWHM) and Maximum Rise Slope vs. Maximum Decay Slope for an Action Potential (MRS vs. MDS) were determined for the different sets in the aforesaid time periods. Recording data visualization, quantification and analysis were done using Clampfit 10.6 software (Molecular Devices) and OriginPro 9.0.0 SR2b87 (OriginLab Corporation). The associated threshold values were determined by matching the first peak of the third derivative with the original trace and the point of intersection was noted as the threshold (Henze and Buzsáki, 2001). The peak action potential amplitude height was measured as a difference between the resting membrane potential of the cell and the peak membrane potential value under evoked responses. The comparison of the number of evoked action potentials at each current injection step was quantified by the total number of action potentials that were observed in one current injection sweep of the current clamp protocol and measured similarly for the entire protocol sweep. The action potential maximum rise and decay slopes, and phase plots were quantified using the rheobase step elicited by our current clamp recording protocol. The FWHM was calculated at the halfway point of the amplitude using Python 3.7.4.

### Statistical Analysis

All data are represented as mean ± SEM for each subset. Normality tests were performed using Kolmogrov-Smirnov Test. Statistical significance analysis for electrophysiological studies was carried out using Mann-Whitney Test for comparative studies with corresponding p values (p < 0.05 considered significant). An unpaired t-test with Welch’s Correction was performed for the comparative statistics of the shaped-based features and inter-event interval of the Ca^2+^ transients using OriginPro 9.0.0 SR2b87 (OriginLab Corporation). Independent t-tests were used for significance analysis for mean firing rates for multielectrode array recordings with p < 0.05 considered significant. The number of asterisks denotes *P < .05, **P < .01, ***P < .001

## ACKNOWLEDGEMENTS

The authors thank T. Gokul Sriman and Rownick Pyne for the technical help and suggestions in regard to the project. We also extend our gratitude to Rosa J. Samuel, Central Animal Facility, IISc Bangalore for the timely supply of rat pups. The research was supported by the Department of Biotechnology, Government of India and the Nanoelectronics Network for Research and Application (NNetRA), Department of Science and Technology, Government of India.

## AUTHOR CONTRIBUTIONS

The project was conceived by SKS and AK. The experiments on multielectrode arrays were performed by SV and analyzed by SV, MR and SG. All electrophysiology experiments were performed by SC and were analyzed by SKS and SC. The two-photon imaging studies were performed by AK and analyzed by SKS and AK. The manuscript was written and edited by SKS, AK, SC, SV, and SG.

## DECLARATION OF INTERESTS

The authors declare no competing interests.

## Notes

### Competing Interest Statement

The authors have declared no competing interest.

